# DRscDB: A single-cell RNA-seq resource for data mining and data comparison across species

**DOI:** 10.1101/2021.01.29.428862

**Authors:** Yanhui Hu, Sudhir Gopal Tattikota, Yifang Liu, Aram Comjean, Yue Gao, Corey Forman, Grace Kim, Jonathan Rodiger, Irene Papatheodorou, Gilberto dos Santos, Stephanie E. Mohr, Norbert Perrimon

**Affiliations:** Department of Genetics, Blavatnik Institute, Harvard Medical School, 77 Avenue Louis Pasteur, Boston, MA 02115, USA; Drosophila RNAi Screening Center, Harvard Medical School, 77 Avenue Louis Pasteur, Boston, MA 02115, USA; European Molecular Biology Laboratory, European Bioinformatics Institute, EMBL-EBI, Wellcome Trust Genome Campus, Hinxton CB10 1SD, UK; The Biological Laboratories, Harvard University, 16 Divinity Avenue, Cambridge, MA 02138, USA; Howard Hughes Medical Institute, 77 Avenue Louis Pasteur, Boston, MA 02115, USA

**Author notes:** Contributed equally (Yanhui Hu, Sudhir Gopal Tattikota, Yifang Liu).

**Keywords:** single-cell RNA-seq, model organisms, data mining, cross-species analysis

## Abstract

With the advent of single-cell RNA sequencing (scRNA-seq) technologies, there has been a spike in studies involving scRNA-seq of several tissues across diverse species including *Drosophila.* Although a few databases exist for users to query genes of interest within the scRNA-seq studies, search tools that enable users to find orthologous genes and their cell type-specific expression patterns across species are limited. Here, we built a new search database, called DRscDB (https://www.flyrnai.org/tools/single_cell/web/) to address this need. DRscDB serves as a comprehensive repository for published scRNA-seq datasets for *Drosophila* and the relevant datasets from human and other model organisms. DRscDB is based on manual curation of *Drosophila* scRNA-seq studies of various tissue types and their corresponding analogous tissues in vertebrates including zebrafish, mouse, and human. Of note, our search database provides most of the literature-derived marker genes, thus preserving the original analysis of the published scRNA-seq datasets. DRscDB serves as a web-based user interface that allows users to mine, utilize and compare gene expression data pertaining to scRNA-seq datasets from the published literature.

## 1. Introduction

Advances in scRNA-seq technologies have enabled a systems-level understanding of several tissues at single-cell resolution across diverse species, resulting in the development of tissue and cell “atlases” [1–3]. Although a vast majority of scRNA-seq studies have been performed in samples from mammals such as mice and humans, a substantial number of studies in less complex model organisms have generated an immense volume of new transcriptomic data at the single-cell level. For instance, in the five years that have followed the availability of microfluidics-based scRNA-seq platform, more than 20 scRNA-seq studies of various organs from *Drosophila,* and across a range of developmental time points and conditions, have been published [4]. Similarly, several other studies on a wide variety of species across the evolutionary tree have been published and scRNA-seq is quickly replacing the more traditional bulk RNA-seq based transcriptomics approach.

The ‘big data’ thus generated from myriad scRNA-seq studies has tremendous potential to aid in the development of algorithms, search tools, and repositories that will benefit the advancement of basic research. Whilst some databases document and compile scRNA-seq studies in one portal (see Supplementary Table 1 for examples), most have caveats that limit their use for cross-species analysis of multiple tissues. These include search databases that focus on one species or tissue and incorporate weak or no ortholog gene search capability. Furthermore, certain databases re-analyze published scRNA-seq data before consuming it. Although re-analysis may not change the transcriptomic architecture of cell clusters, it may change the structure of the scRNA-seq map and the set of top-enriched marker genes as compared with the original analysis published by the authors. Moreover, search databases that feature scRNA-seq datasets from multiple species to facilitate a cross-species survey of a given gene tend to work well for orthologous genes that have the same name (or keyword) in different species but might not map orthologs with different names. Altogether, there is a need for comprehensive databases that allow users to search genes of interest across various scRNA-seq datasets obtained from different species and preserve the outcomes of the original published analyses.

To provide a comprehensive scRNA-seq data mining resource with the features discussed above, we developed DRscDB (***D**rosophila **R**NAi Screening Center’s **s**ingle-**c**ell **D**ataBase*), which provides a simple, user-friendly web search tool. DRscDB allows users to search a gene of interest and identify not only all of the relevant datasets expressing this gene but also all of the datasets in which an orthologous gene from another species is expressed. Information about relevant tissues and cell types is summarized at DRscDB in an easy-to-read table and can be visualized as dot plots, heatmaps or bar graphs. In addition, DRscDB can also facilitate enrichment analysis of a user-provided list of genes. Specifically, genes on the list are compared to top markers of various cell types from all datasets across species and the best-matched cell clusters of relevant tissues are returned. This function can help users assign clusters when a new scRNA-seq dataset is obtained and can also help users identify the most relevant cell type/tissues if a gene list is obtained from another source, such as a cell-based screen or functional annotation. Altogether, DRscDB makes it possible for users to mine multiple scRNA-seq datasets with ease and is unique in its design and use of a robust cross-species ortholog search feature (Supplementary Table 1).

## 2. Material and methods

### 2.1. Standardization of information from the literature

Most scRNA-seq datasets are analyzed and presented differently in different journals, making it challenging to fetch marker genes in an automated manner. Hence, we decided to manually curate published scRNA-seq papers. To optimize the choice of papers and their metadata for manual curation, we streamlined our approach based on two major criteria. First, we focused on a defined set of species and second, we sought to cover most major tissue types common among these species. To achieve optimal coverage of evolutionarily conserved model organisms ranging from small invertebrates to larger vertebrates, we decided to include scRNA-seq studies of tissues from *Drosophila melanogaster* (fruit fly), *Danio rerio* fzebrafish), *Mus musculus* (mouse), and *Homo sapiens* (human). Next, we focused on major tissue types that were analyzed in *Drosophila* and whose analogous tissues had been studied in the other species. Applying these criteria, we searched the published literature and manually curated relevant scRNA-seq papers. The required metadata, which includes the lists of marker genes presented by the authors in the respective papers, were collected and stored in Excel files of a standard format along with supporting information specific to each dataset or study. The metadata and marker gene files include all of the essential details such as paper title, brief summary, species and tissue type, type of scRNA-seq method, number of clusters identified, and, importantly, the list of marker genes along with statistics regarding differential gene calling, such as fold enrichment and P values if available. In cases where information pertaining to the marker genes were not reported in the supplementary information, we collected the marker genes per cluster that were discussed in the main text of the paper.

### 2.2. Data processing pipeline

To understand whether a gene is expressed in each cell type, as well as the expression scale and level, additional data files, specifically the raw gene-to-cell expression matrix and metadata file, are needed as input files for the data processing pipeline. The gene-to-cell expression matrix contains the raw unique molecular identifier (UMI) counts or read counts, and the metadata file contains barcode and cell type annotation information. For most publications selected, the matrix and metadata files were available from GEO records associated with the manuscript. In cases where such files were not deposited in public repositories, we requested the files from the corresponding author. Next, the matrix and metadata files were processed using a customized script based on Seurat [5]. The output includes two files, clusterMetadataTable and gene2clusterTable. The clusterMetadataTable file is a summary at cluster level generated from metadata file. The gene2clusterTable file is generated by extracting the data from the Seurat DotPlot function and contains information about the percentage of cells expressing and average expression levels of each gene in each cluster (Supplementary Fig. 1). The data processing pipeline can significantly compress the data. For example, the file size of the gene2cell expression matrix of the *Drosophila* blood dataset from [6] is about 500 MB, but after processing through this pipeline, the output file for the gene2cluster matrix is about 10 MB. All of the scripts used for the data processing pipeline for each publication are stored in GitHub and are available to public (https://github.com/moontreegy/scseq_data_formatting).

### 2.3. Data import and synchronization

The gene identifiers used for gene markers and data matrices from different publications vary in terms of both the source and the version. We synchronized gene identifiers to the current NCBI Entrez GeneID and species-specific identifiers such as Mouse Genome Informatics (MGI) gene identifiers for mouse genes and FlyBase gene identifiers for *Drosophila* genes. Identifiers that cannot be mapped are excluded. Gene identifiers will be updated periodically to reflect updates at FlyBase and other resources.

### 2.4. Marker gene selection, ortholog mapping and enrichment analysis

The marker genes, as well as statistics such as fold enrichment, P values and/or adjusted P values, were usually collected during curation process from either supplemental file associated with the publication or GEO. Very often hundreds and sometimes a few thousands of marker genes were identified in each cluster and full marker gene lists are reported in all but a small number of publications that only make available the subset of marker genes that were validated. When there are more than 100 marker genes reported per cluster, the top 100 marker genes per cluster are selected based on the fold change, using an adjusted P value or a P value of 0.05 as a filter. The gene components of each gene set are then mapped to other model organisms using DRSC Integrative Ortholog Prediction Tool (DIOPT) vs8 with a filter applied to map only high or moderate rank results [7]. Enrichment P value of marker gene sets is calculated based on the hypergeometric distribution. The enrichment strength can be illustrated using either the negative log10 of P value or the fold change at DRscDB website.

### 2.5. Implementation of the online resource

The DRscDB web tool (https://www.flyrnai.org/tools/single_cell/) can be accessed directly or found at the ‘Tools Overview’ page at the DRSC/TRiP Functional Genomics Resources website (https://fgr.hms.harvard.edu/) [8]. The back end was written using PHP with the Symfony framework. The front-end HTML pages take advantage of the Twig template engine. The JQuery JavaScript library with the DataTables plugin is used for handling Ajax calls and displaying table views. D3 and Vega-Lite javascript libraries are used for the graphing (http://idl.cs.washington.edu/papers/vega-lite/). The Bootstrap CSS framework and some custom CSS are also used on the user interface. A mySQL database is used to store the curated information as well as cluster-level statistics. Both the website and databases are hosted on the O2 high-performance computing cluster made available by the Research Computing group at Harvard Medical School.

## 3. Results

### 3.1. Literature curation and data process

Single-cell RNA-sequencing (scRNA-seq) provides a powerful way to study transcriptomes at single-cell level, allowing researchers to uncover potential new biological insights not discoverable using traditional bulk RNA-seq methods. For example, analysis of scRNA-seq data can reveal rare cell populations, uncover regulatory relationships between genes and cells, and be used to trace the trajectories of various cell lineages [9]. The number of publications reporting scRNA-seq data is increasing dramatically and thus far includes more than 20 publications for *Drosophila melanogaster. \Ne* selected 18 publications with scRNA-seq datasets for *Drosophila* [4] as well as 29 publications for human, mouse, zebrafish for our initial manual curation efforts. Information such as tissue, developmental stage, sequencing method, cell type, and marker genes were collected from original publications and reorganized using a standard template file. By design, we use the marker gene lists provided by the papers and do not reanalyze the datasets, thus preserving the results of the original analyses done by researchers with specific relevant expertise. In addition, the data files that had been deposited by study authors into NCBI GEO were retrieved and processed to obtain cluster-level statistics such as the percentage of cells that express a given gene and average expression levels for the gene within a cluster. We found that for 36% of the publications (17 of 47), the information publicly available at the time of our curation was not adequate for data processing. For example, some data files were missing an annotation file of cell barcodes used to cluster the cells, and/or missing a cell-gene expression matrix. For publications in this category, we requested the additional information from the corresponding authors and in several cases (4 publications) the information was kindly provided by the authors. When we were unable to obtain the data, we only imported the marker genes as extracted from the publication (Fig. 1). Standardized information was then imported into a mySQL database. During import, gene symbols and/or identifiers were mapped and synchronized to common identifiers (Entrez GeneIDs) and/or species-specific identifiers (e.g. FlyBase FBgn IDs for *Drosophila* genes, MGI IDs for mouse genes). In addition, the top 100 marker genes were selected and mapped to orthologous genes of the other species represented in the database using DIOPT (Fig. 1) [7].

**Figure 1.**
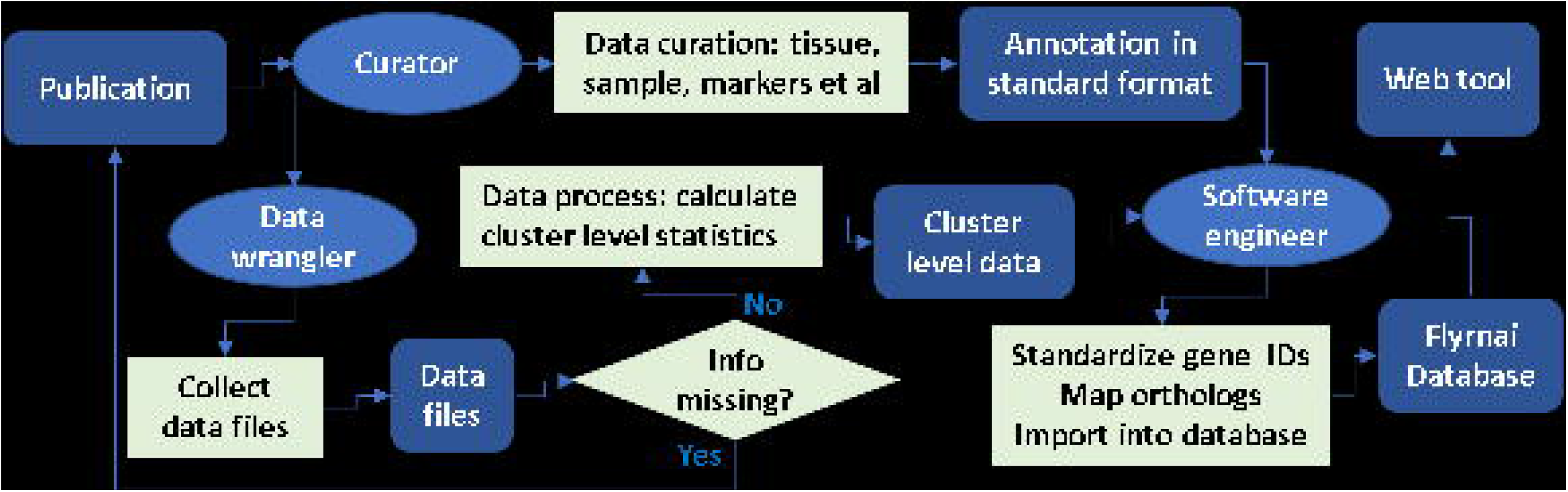
Curation and processing of scRNA-seq datasets from the literature. DRscDB is built based on the curation effort of published scRNA-seq literature. During the curation process, curators extract the information about experimental design, sample information as well as marker genes from each publication and organized the information in standard template while data wranglers retrieve the data files (cell expressing matrix and metadata file) from GEO and calculate the expression statistics of each gene at cluster level (Supplementary Fig. 1). Subsequently, data files and annotation files are processed by a software engineer for database upload.

In summary, basic information about experimental design, sequencing platform, and analysis tool used, and analysis results associated with the scRNA-seq datasets, such as cell types and marker genes, are captured during the curation stage. Further, detailed information regarding the percentage of cells that express a given gene and average expression levels for a gene within a cluster are captured during a data processing stage. Curated information and processed data are synchronized and stored in the database. At the same time, the top marker genes are selected and mapped to orthologous genes for enrichment analysis.

### 3.2. DRscDB as an online resource

DRscDB is an online resource built to access the curated information and processed data from scRNA-seq publications. When multiple sample types were sequenced, e.g. different treatment conditions, genotypes, sex, or developmental stages, the datasets were processed separately, allowing users to compare gene expression changes across different samples. For example, the publication from Tattikota et al. (2020) includes the scRNA-seq datasets for *Drosophila* hemocytes from unwounded control flies, wounded flies, and wasp-infested fly samples [6], and we processed each dataset separately as well as jointly, thereby importing 4 datasets into DRscDB. Upon mining the data in DRscDB, users can easily compare expression patterns across samples. Currently, DRscDB covers 90 datasets from 47 publications (Table 1, Supplemental Table 2).

**Table 1.** DRscDB coverage

| Species | Tissue | # publications | # datasets | PMIDs |
| --- | --- | --- | --- | --- |
| Drosophila | Immune | 4 | 12 | 32396065[6];32900993[10];32162708[12];32487456[11] |
| Drosophila | Ovary | 3 | 4 | 31919193[13];32339165[14];33159074[15] |
| Drosophila | Wing disc | 3 | 5 | 31363221[19];32815271[20];31455604[21] |
| Drosophila | Brain | 2 | 11 | 31746739[22];29909982[23] |
| Drosophila | Intestine | 2 | 2 | 31851941[24];31915294[16] |
| Drosophila | Embryo | 1 | 1 | 28860209[25] |
| Drosophila | eye disc | 1 | 4 | 30479347[26] |
| Drosophila | Kidney | 1 | 1 | 32175841[27] |
| Drosophila | Testis | 1 | 1 | 31418408[28] |
| Human | Kidney | 5 | 14 | 29870722[29];31506348[30];29980650[31];30789893[32];NA[33] |
| Human | Brain | 2 | 3 | 31303374[34];31042697[35] |
| Human | Immune | 2 | 2 | 28428369[36];31413257[37] |
| Human | Intestine | 2 | 5 | 31753849[18];30526881[38] |
| Human | Testis | 2 | 5 | 30726734[39];30315278[40] |
| Mosquito | Immune | 1 | 2 | 32855340[17] |
| Mouse | Kidney | 4 | 4 | 31689386[41];29794128[42];31118232[43];29622724[44] |
| Mouse | Testis | 2 | 2 | 31237565[45];29695820[46] |
| Mouse | Brain | 1 | 1 | 29545511[47] |
| Mouse | Embryo | 1 | 1 | 30840884[48] |
| Mouse | Immune | 1 | 1 | 27365425[49] |
| Mouse | Intestine | 1 | 1 | 29144463[50] |
| Zebrafish | Brain | 3 | 6 | 29608178[51];31018142[52];29576475[53] |
| Zebrafish | Immune/Kidney | 1 | 1 | 28878000[54] |
| Zebrafish | Intestine | 1 | 1 | 32092251[55] |

DRscDB allows users to search a gene of interest using a ‘single gene search page’. In this case, the input gene and its predicted orthologs are retrieved along with the number of datasets in which each gene was found to be expressed. For example, a search with the human gene *GAD1* returns a table including the *GAD1* gene and orthologs such as *Gad1* in mouse and fly, and *gad1a* and *gad1b* in zebrafish. There are 24 human datasets in which *GAD1* is annotated as expressed, 8 datasets for the mouse ortholog, and 6/7 datasets for each of the zebrafish orthologs, and 28 datasets for fly *Gad1.* This information is summarized in an additional column along with orthologous gene search results. Users have the option to view detailed information about relevant scRNA-seq datasets for any of the genes in the summary table. The results of data mining of tissues and cell types that express a gene of interest are first presented in a summary table. Users can also view cluster-level statistics as a dot plot, bar graph, or heatmap. With a dot plot, the percentage of cells that express the gene is represented by the size of each dot and the average level of expression is represented by the intensity of the color. Users can quickly survey the level and scale of cells expressing a gene of interest in all relevant cell types and across all tissues, based on the publications covered in the database. In the summary table, information about whether a gene of interest is a cell-type specific marker is included, and users also can view marker statistics, which are presented as bar graphs in which the height of the bar represents the fold enrichment of gene expression in a cluster as compared to the rest of the cells, and the color intensity of the bar represents the enrichment P value (Fig. 2).

**Figure 2.**
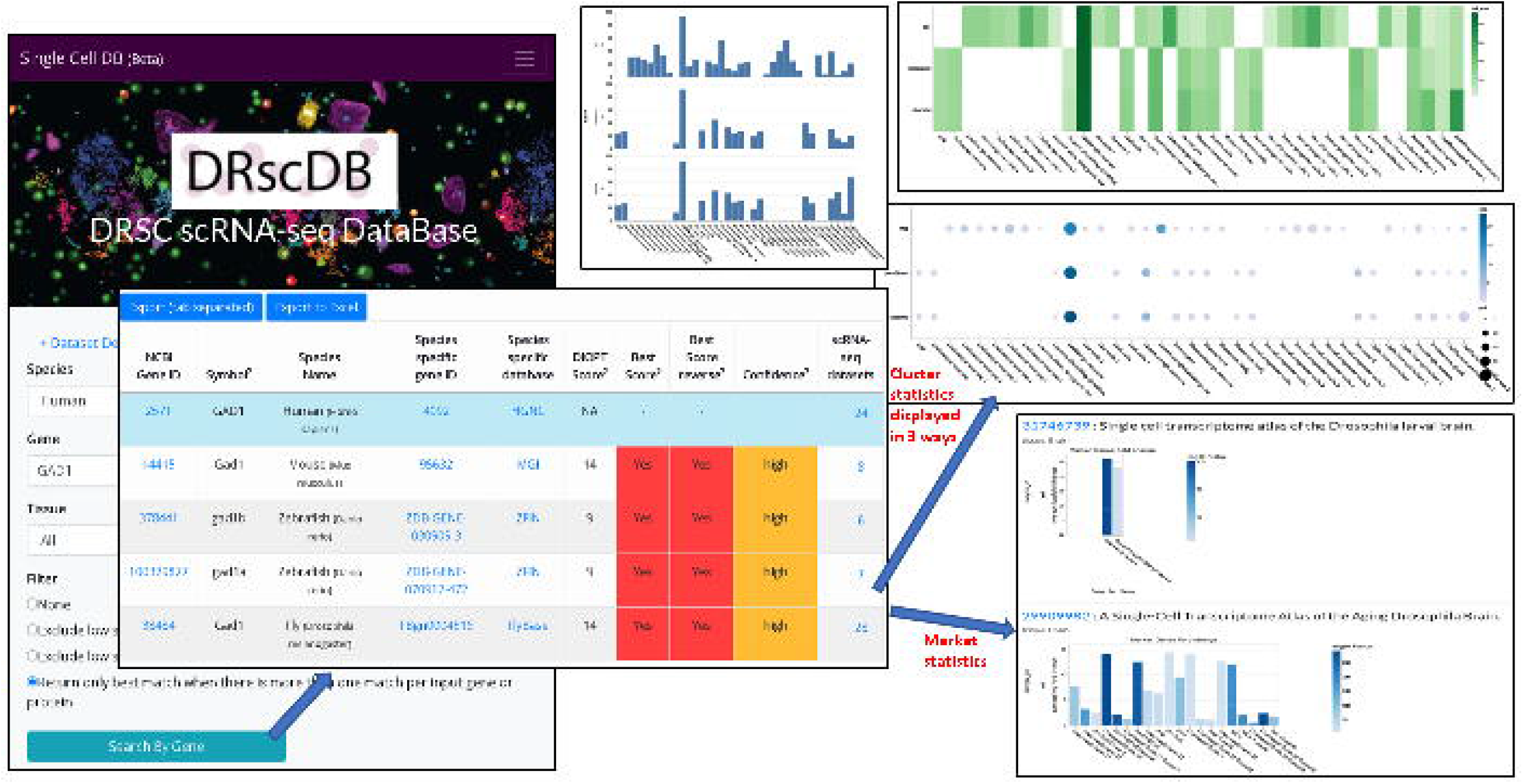
Use of DRscDB for data mining. At the DRscDB search page, a user can enter the gene of interest with or without specifying the tissue of interest, the result is summarized in a table format listing the number of datasets expressing the gene of interest as well as the orthologous genes. Then the user can find more detailed information such as the relevant clusters expressing the gene of interest. The statistics about the percent of cells expressing the gene as well as the average expression level can be visualized by dot plot, bar graph, or heatmap. If a gene is identified as one of the marker genes for any of the clusters, the statistics of fold enrichment as well as P value are also displayed by bar graph.

We selected up to 100 top markers for each cluster based on log2 fold change (log2fc) as well as the adjusted P value or P value to generate the gene sets used for enrichment analysis. In addition, all of the gene sets are also mapped to other model organisms using DIOPT. When a user enters a list of genes at the enrichment page, the input genes are compared with the gene sets built based on the top markers from all datasets, and the most similar gene sets are returned in a display that includes statistics, e.g., fold change and P values. Users also have the option to enter multiple gene lists, using the first column to specify the name of the list and the second column to specify the gene components of each list. This function allows users to compare the gene markers for all clusters from one scRNA-seq study with the gene markers from a different study from the same or a different species. This then allows users to identify similar clusters as defined by top markers. The results can be visualized at DRscDB as heatmap or a dot plot (Fig. 3, Supplementary Fig. 3).

**Figure 3.**
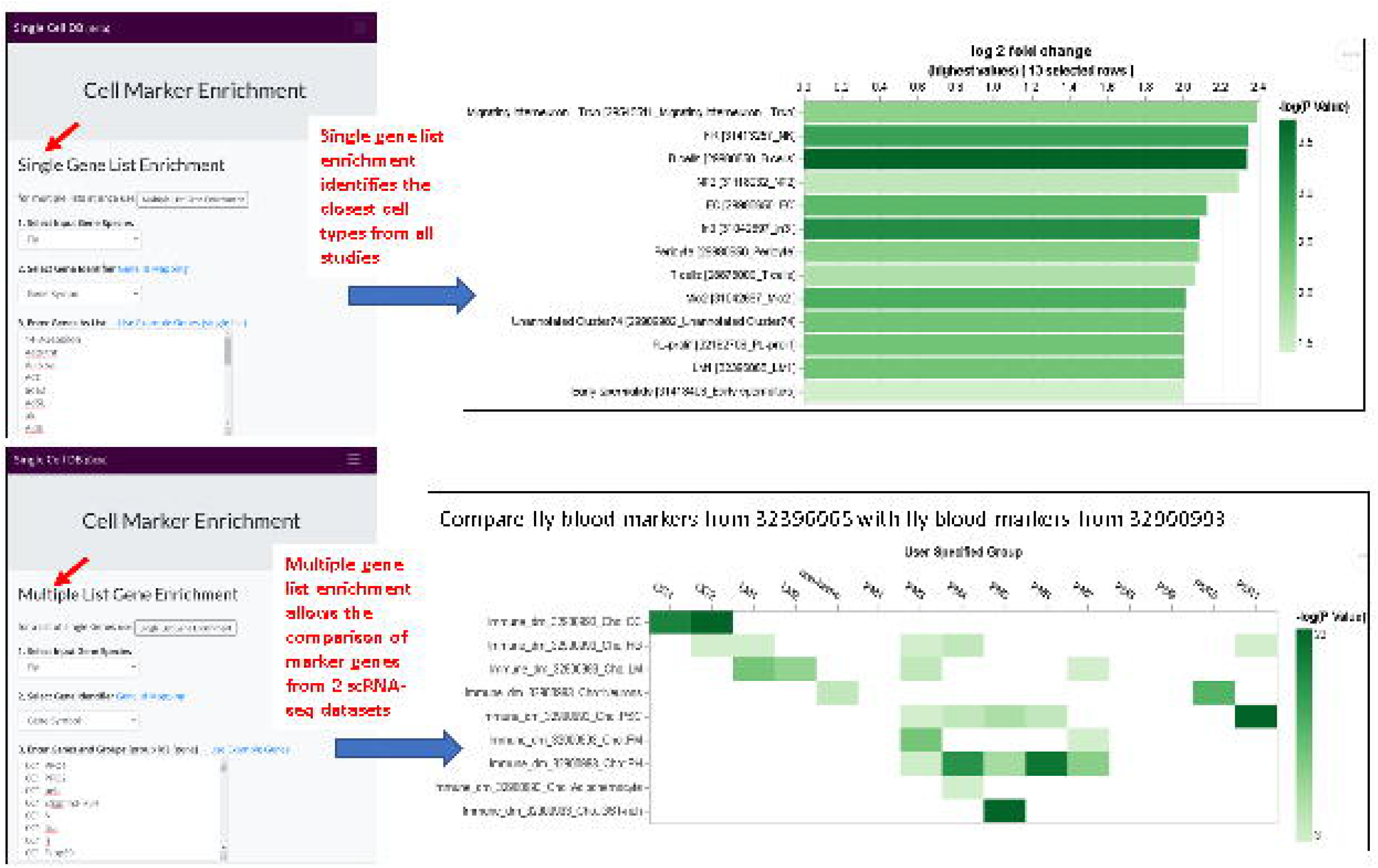
Use of DRscDB for enrichment analysis. At the DRscDB enrichment analysis page, a user can input a list of genes and find out the clusters for which the input genes are significantly enriched among the top 100 marker genes. In addition, at this page, a user can also enter multiple gene lists and compare each input gene list (for example, 15 lists) with every cluster of a selected study (for example, 9 clusters). The enrichment results are visualized by a heatmap, consisting in this example of a 9×15 matrix, with columns representing each input gene list and rows represents each cluster from the selected study. The darkness of the color represents similarity (-log10 P value or fold enrichment).

Given the exponential increase in publications that include scRNA-seq datasets, it is not practical for one group to manually curate and annotate all publications. Hence, to further increase the coverage of DRscDB, we implemented a webpage that makes it possible for community members to submit information corresponding to studies or publications currently not covered in DRscDB. To do this, the submitter would fill in information in a set of template curation files. Participation in this researcher-driven submission process will expedite our internal paper curation and import process. At DRscDB website under the menu option “Datasets”, a summary page of all the publications imported so far is dynamically generated from the database, which allows the researchers to check if a paper has been included (https://www.flyrnai.org/tools/single_cell/web/summary).

### 3.3. Applications of DRscDB

Currently, *Drosophila* single-cell RNA-seq datasets cover a wide range of tissues, including the brain, wing disc, intestine, immune or hematopoietic cells, and reproductive system, some of which are also covered by multiple datasets (Table 1). In particular, there are four published datasets for *Drosophila* myeloid-like immune cells and 48 cell clusters in total [6, 10–12]. We systematically compared the top markers (up to 100) of each cluster with the top markers of every other cluster from these four publications and calculated a similarity score for each comparison, i.e., the negative log10 of enrichment P value (-log10 P value). Then, we built a 48 x 48 similarity matrix and performed unsupervised hierarchical clustering. The clustering results show that blood cell types such as plasmatocytes, crystal cells, and lamellocytes from different studies align quite well with each other (Fig. 4), indicating that our mapping algorithm works efficiently. Next. we executed a similar strategy with another tissue, the ovary, with three published scRNA-seq datasets [13–15]. Comparison of the top markers among all the cell clusters from these three studies also show that most similar cell types, including germline stem cells, align and cluster together, with very high similarity scores (Supplementary Fig. 2).

**Figure 4.**
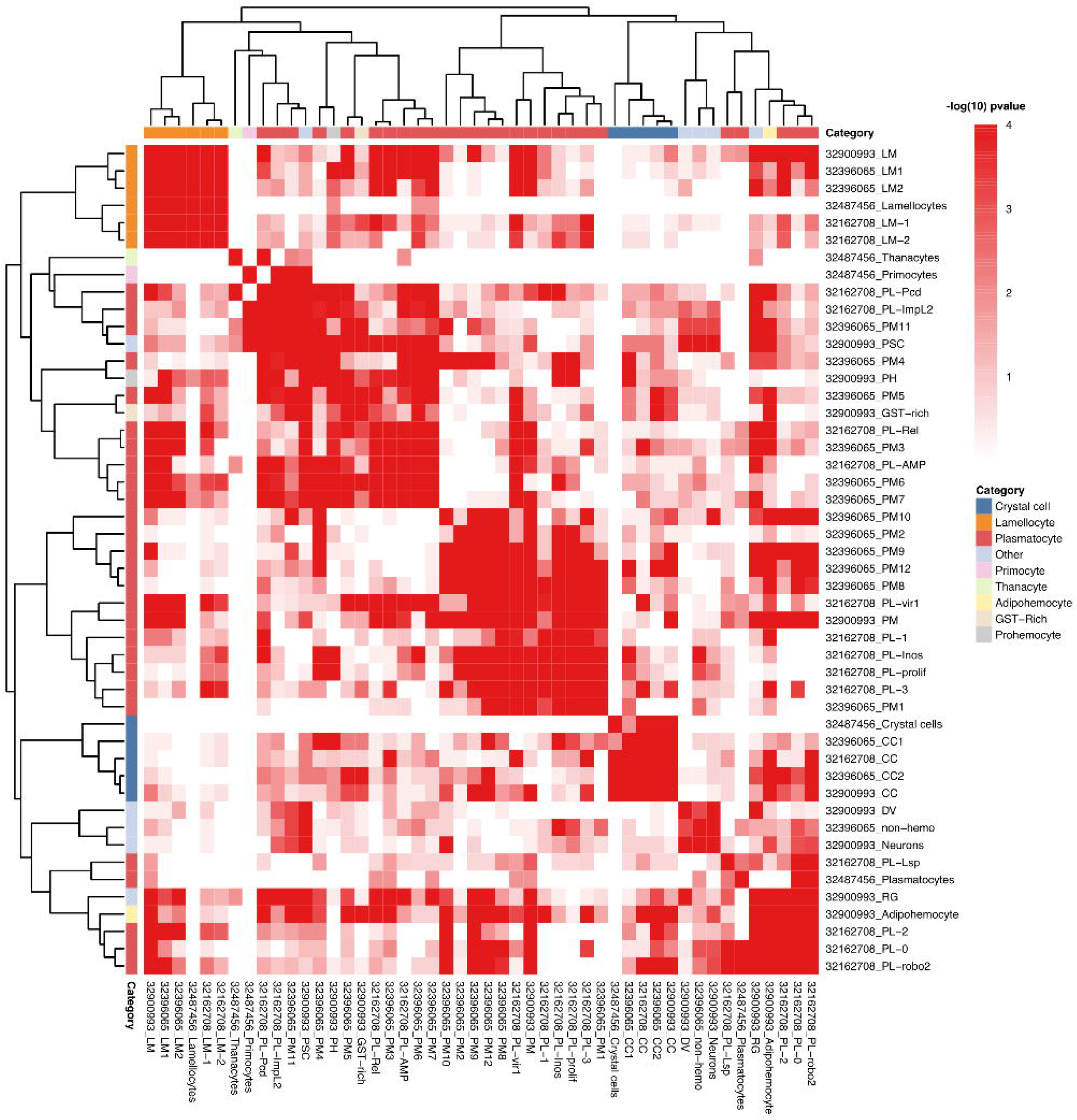
Unsupervised hierarchical clustering of enrichment results comparing top markers from for publications on the *Drosophila* immune system. Clustering of the top 100 marker genes per cluster from [6], with 3 other published immune datasets [10–12]. The results reveal that similar cell types tend to cluster together from these four immune datasets; therefore, it is reasonable to suggest that for newly generated datasets DRscDB can be used to assign cell types.

To further test whether the literature-derived marker genes can be used to associate cell clusters from different datasets, as well as test the comparison across species, we started with two scRNA-seq datasets pertaining to *Drosophila* blood and intestine [6, 16]. First, to facilitate a query of cluster groups and their respective marker genes, we introduced a gene set query feature that allows users to simply copy and paste “cluster_id, gene symbol” into the query box and choose a published dataset to match query cell clusters. In order to demonstrate if cluster to cluster matching effectively works, we simply compared the top ten marker genes of *Drosophila* blood dataset with all the marker genes of the same dataset [6] to test if each cluster maps to its own cluster. As expected, each cluster significantly overlapped with its own cluster (Supplementary Fig. 3a). Next, we performed a similar task with the *Drosophila* intestine dataset [16] and observed a similar tight cluster to cluster correlation (Supplementary Fig. 3b). These results show that DRscDB can effectively perform a one-to-one correlation analysis of query clusters available in various datasets.

Next, to demonstrate the power of this feature across species, we compared *Drosophila* blood marker genes [6] with marker genes derived from A. *gambiae* (mosquito) blood scRNA-seq [17]. Of note, *Drosophila* blood cluster ‘PM2’ significantly correlated to ‘HC4’ in mosquito blood, both of which have been described as proliferating hemocytes that express mitotic marker genes [6,17]. Likewise, ‘PM6’ in flies mapped well to HC6 in mosquitoes, and both clusters are enriched in genes that encode antimicrobial peptides (AMP). In addition, the ‘HC1’ cluster identified in the mosquito blood study maps to mature crystal cell cluster ‘CC2’, and both express high mRNA levels of prophenoloxidases (PPO), which are characteristics of crystal cells or otherwise called oenocytoids [6, 17]. Interestingly, ‘HC1’ also highly correlated with ‘PM8’, a plasmatocyte cluster that is enriched in peroxiredoxins. Surprisingly, the mosquito fat body (FBC) correlated with fly ‘LM1’, a potential lamellocyte intermediate cluster, and the mosquito muscle clusters (MusC) correlated with ‘PM6-7’, which represents PM^AMP^ (Fig. 5A). We also queried the *Drosophila* intestine dataset [16] against its human counterpart [18]. As expected, the fly intestinal cell clusters representing enterocytes, intestinal stem cells and enteroblasts (ISCs and EBs), and enteroendocrine cells significantly overlapped with the analogous human cell types: enterocytes, stem/progenitor cells and transiently amplifying cells, and enteroendocrine cells, respectively (Fig. 5B).

**Figure 5.**
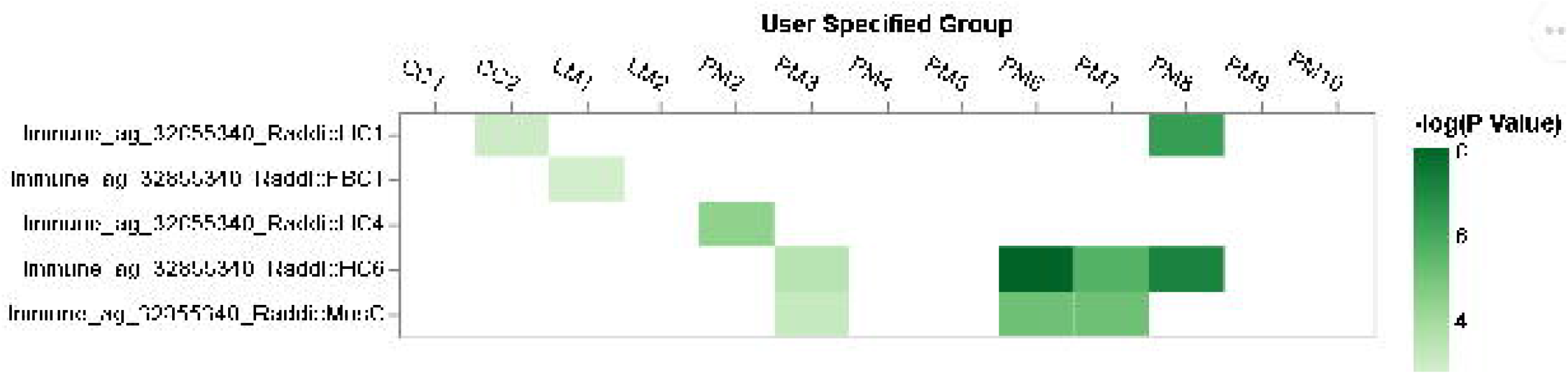
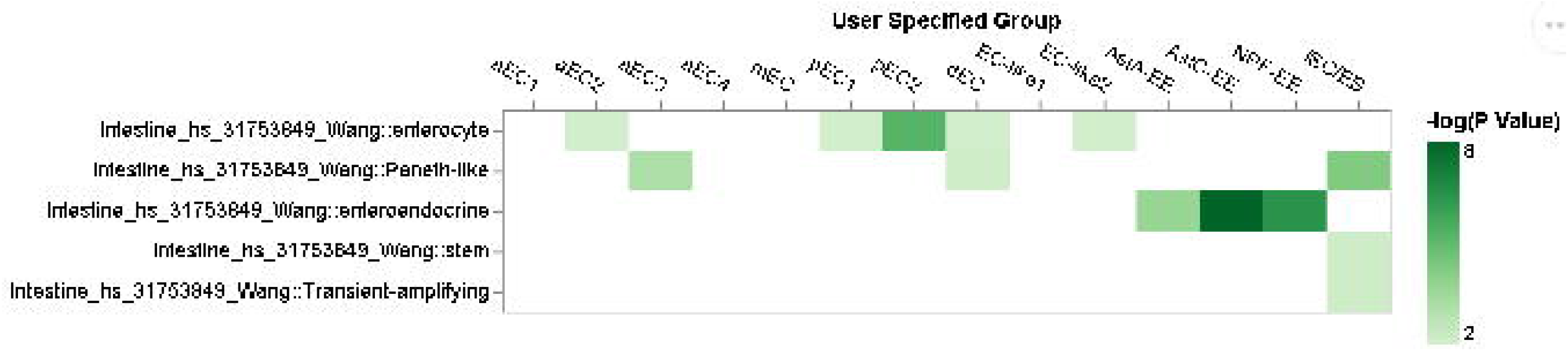
DRscDB facilitates comparison of cell clusters across datasets and species. A. Comparison of the top 10 marker genes per cluster derived from *Drosophila* [6] or mosquito [17] blood scRNA-seq datasets. B. Comparison of the top 20 marker genes per cluster from Hung et al., 2020 [16] with published human intestinal cell clusters [18].

Altogether, DRscDB can efficiently match cell clusters in a query to those from the published literature from different datasets and species, thereby allowing for cross-dataset and cross-species comparisons of cell clusters. When a new scRNA-seq dataset is obtained, researchers typically review the marker genes for each cluster and as much as possible, assign cell types based on prior knowledge. DRscDB facilitates comparison of marker genes from a new dataset to marker genes from any relevant publications included in the database. Thus, the analyses supported by DRscDB can help facilitate the process of cell type assignment.

## 4. Discussion

### 4.1. Querying genes across datasets and species

DRscDB standardizes and centralizes scRNA-seq datasets for *Drosophila,* human, and other common genetic model organisms, making it possible for users to mine scRNA-seq data and perform cross-species analyses easily. DRscDB is unique in that it uses the powerful DIOPT approach to map orthologous genes, such that it is able to facilitate efficient and robust gene searches across species. Furthermore, DRscDB’s cell type enrichment tool allows for systematic matching and categorization of similar cell clusters identified in various datasets.

DRscDB was also designed to use information taken from original publications, thus leveraging the biological expertise of the submitting scientists and providing a centralized resource that is complementary to the consortium efforts such as the Human Cell Atlas (HCA), Mouse Cell Atlas (MCA), Fly Cell Atlas (FCA) and Single Cell Expression Atlas at EMBL-EBI, which rely on reanalysis of raw datasets. One reason this is helpful is because although reanalysis of published scRNA-seq datasets provides an unbiased perspective of the published data, it may not replicate results that can only be obtained by detailed manual analysis. As a result, the structure of the scRNA-seq maps and top enriched genes in these repositories may differ from those reported in the original analyses. Nevertheless, as we recognize the value of consortium efforts and re-analyzed data, DRscDB was designed with the flexibility to receive data and annotations directly from centralized resources in the future. Thus, DRscDB preserves and integrates scRNA-seq analyses available in published literature, and serves as a flexible and potentially comprehensive online resource.

### 4.2. Toward the identification of evolutionarily conserved cell populations

DRscDB facilitates efficient cell cluster identity and the identification of its potential match in other species. We demonstrated its use by querying cell clusters in DRscDB with the top marker genes from the studies as the user input, for both Drosophila blood and intestine datasets, and further compared these with the analogous tissues in mosquito or human, respectively. Although other such ‘cell assign’ tools are available and help in revealing the potential true identity of a particular cell cluster in question, most of these tools have been developed for mammalian tissues. To our knowledge, tools that allow for cross-species ‘matchmaking’ among query and published cell clusters are limited. Hence, DRscDB stands out by allowing users to quickly understand the relationships between new cell clusters or clusters of interest in one species and clusters from other samples and species available at DRscDB. We do note, however, that for certain tissues, such as *Drosophila* blood, most clusters could not be mapped efficiently to blood datasets of other vertebrate species. We believe that criteria such as the number of marker genes and/or additional published datasets may help to streamline cluster matching across diverse species.

### 4.3. Community participation and future direction

DRscDB has the potential to be a rich resource of published scRNA-seq datasets that can be of high practical value to researchers, students, teachers, and others. As mentioned, DRscDB relies on manual curation of the published literature as a means of preserving the original analyses done by the study authors. Although manual curation has benefits, it also suffers from being time consuming and labor intensive, such that inclusion of some published datasets might be delayed or unsupportable. With this in mind, we provide an option that allows those conducting scRNA-seq studies to upload relevant information, thus facilitating the addition of more datasets, which will further enhance ability to search across species as supported by DRscDB query tools. Thus, we hope that by including a mechanism for community annotation, the coverage and usefulness of the resource will continue to expand.

## Supporting information

legend for supplement figures and tables

Supplementary figur2

## Availability

Code and results from data processing are located at https://github.com/moontreegy/scseq_data_formatting. The online resource is available without restriction at https://www.flvrnai.org/tools/single_cell/web/.

## Acknowledgements

We would like to thank the members of Perrimon laboratory, FlyBase consortium, Drosophila RNAi Screening Center (DRSC) and Transgenic RNAi Project (TRiP) for helpful suggestions regarding the project. We would like to thank Drs. A. McMahon, M. Ariss and A. Ghosh for their input and suggestions during the development of DRscDB. Finally, we are immensely thankful to Drs. L. Zhao, S. Aerts, S. Sprecher, R. Xi, R. Lehmann, D. Geschwind, A. Elkins, M. Deng, B. Humphrey, D. Conrad, M. Boutros, A. Teleman, M. Zappia, M. Frolov, G. Raddi, O. Billker, and C. Barillas-Mury for kindly sharing additional data files pertaining to their respective scRNA-seq studies. This work was supported by NIH NIGMS P41 GM132087. N.P. is an investigator of Howard Hughes Medical Institute.

