## Supplementary material for "DRscDB: A single-cell RNA-seq resource for data mining and data comparison across species": legend for supplement figures and tables

**Supplementary Figures**

**Supplementary Figure 1. Overview of the data processing pipeline**

The cell2cluster annotation and gene2cell expression matrix are retrieved from GEO. Then, the annotation and matrix are further processed and analyzed by our customized script. Finally, the cluster metadata table and gene2cluster matrix are integrated into DRscDB.

**Supplementary Figure 2. Unsupervised hierarchical clustering of enrichment result comparing top markers from 3 publications of Drosophila ovary**

Clustering of the top 100 marker genes per cluster from three published fly ovary datasets [13-15]. The results show that similar cell types from these three ovary datasets tend to cluster together.

**Supplementary Figure 3. DRscDB facilitates comparison of cell clusters across datasets and species**

A. Querying the top 10 marker genes per cluster from Tattikota et al., 2020 against the top100 marker gene sets pertaining to the same study resulted in a significant correlation between each cluster pair. B.Querying the top 10 marker genes per cluster from Hung et al., 2020 against the top100 marker gene sets pertaining to the same study resulted in a significant correlation between each cluster pair.

**Supplementary Table 1: Review of existing resources**

**Supplementary Table 2: Curated information about each publication**
