## Supplementary figures and images for "DRscDB: A single-cell RNA-seq resource for data mining and data comparison across species"

### sfigure1.png

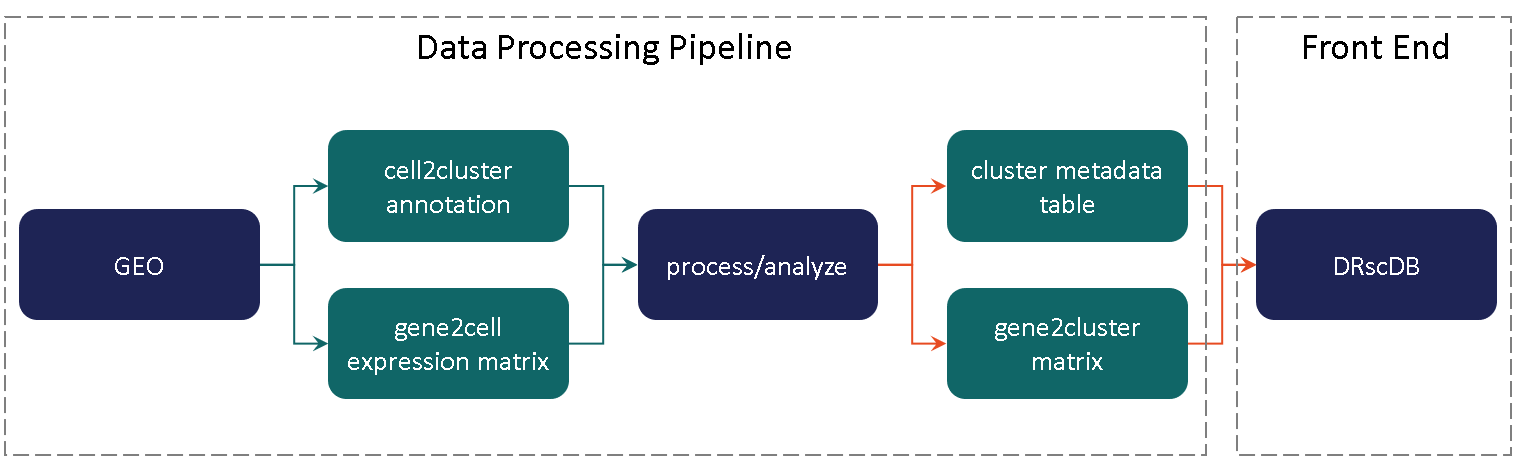

### sfigure2.png

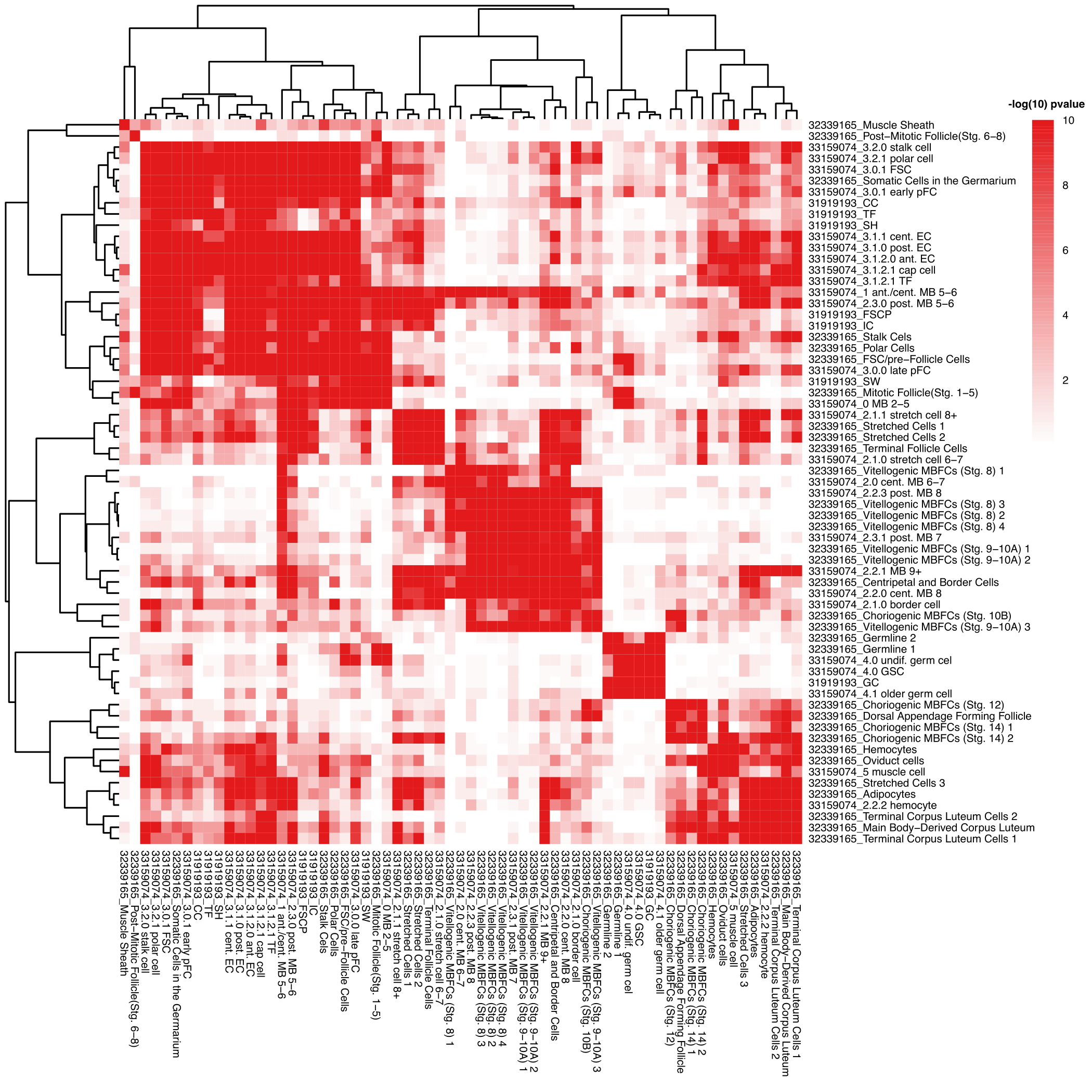

### sfigure3A_final.png

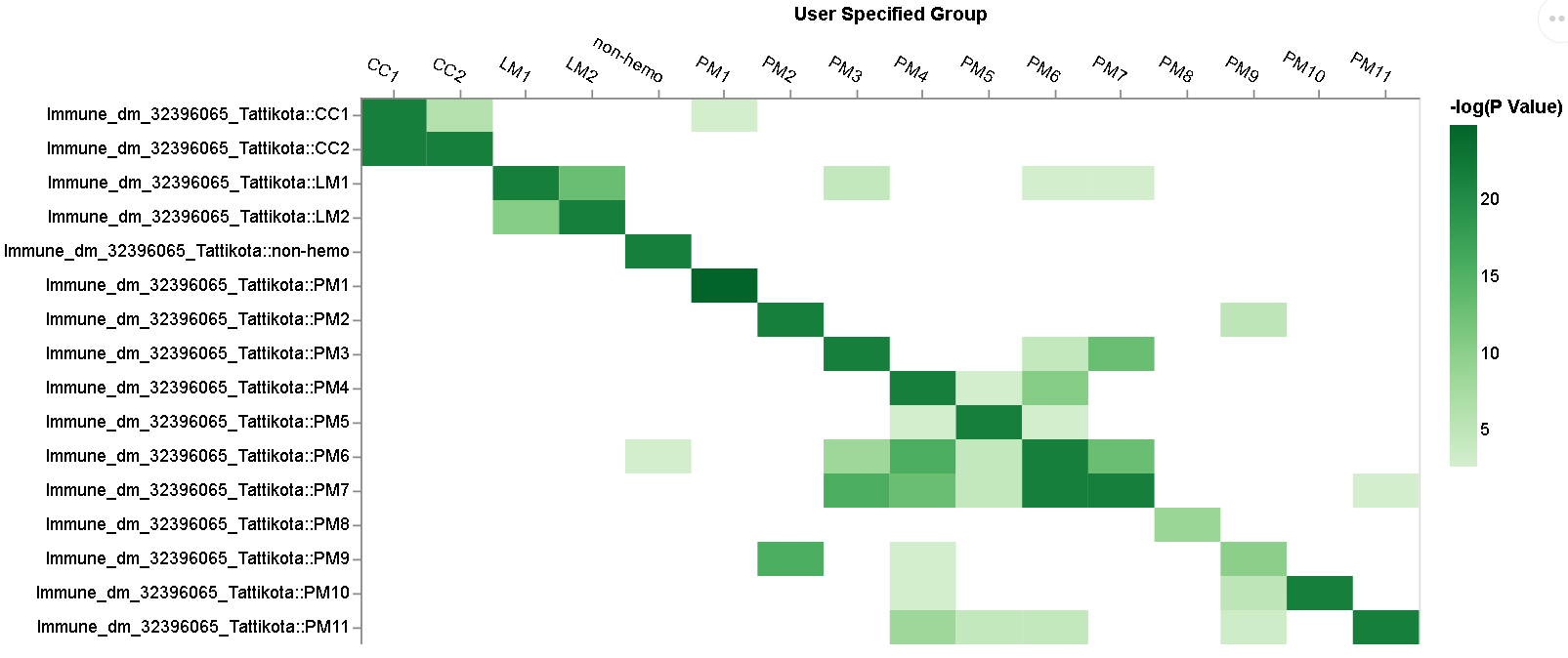

### sfigure3B_final.png

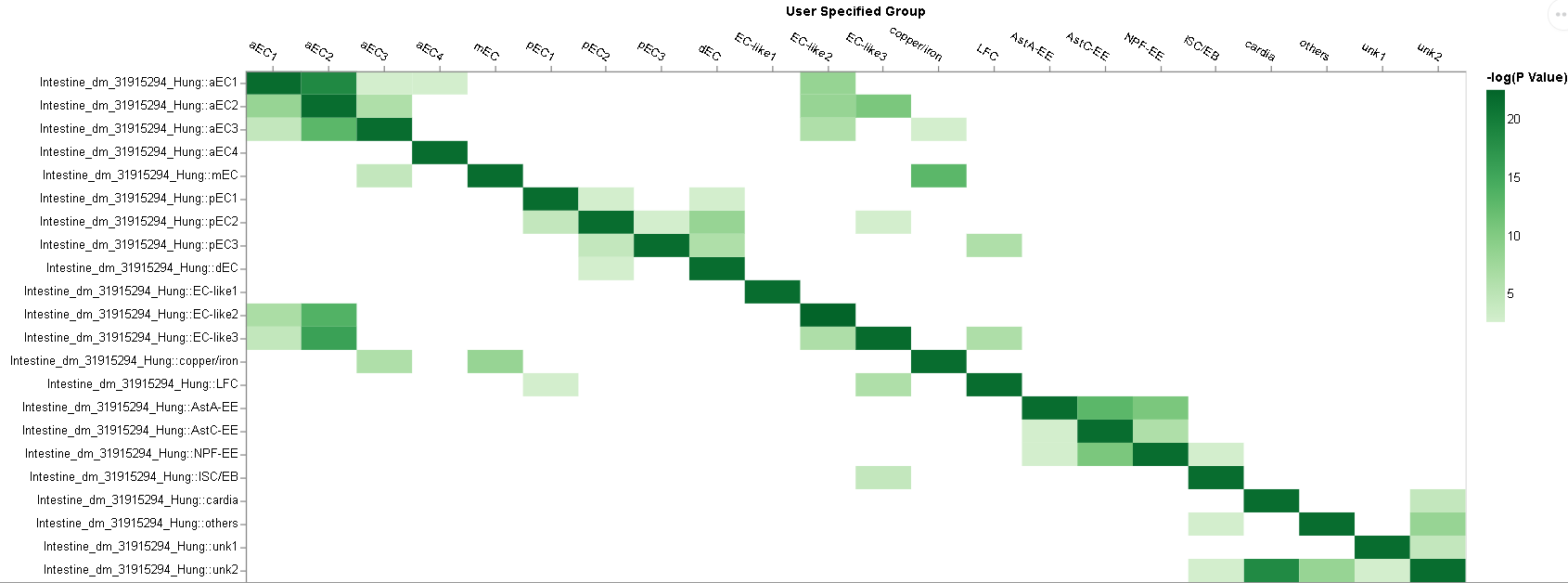
